## Supplementary Data for "Integrative *in vivo* analysis of the ethanolamine utilization bacterial microcompartment in *Escherichia coli.*"

### Supplementary tables

| Primer name | Sequence |
|---|---|
| F_bb1 | TGAGCTAGCTGTCAAGGATCC |
| R_bb1 | AAGTTAAAATAAGGCTAGTCCGTTAT |
| F_eutGgRNA | ATCCTTGACAGCTAGCTCAGTCCTAGGTATAATACTAGTCGGCACACCTTCGGTCAATG |
| R_eutGgRNA | ACTAGCCTTATTTTAACTTGCTATTTCTAGCTCTAAAACCATTGACCGAAGGTGTGCCG |
| F_bb2 | CATAGTTAAGCCAGCCCCGA |
| R_bb2 | TGTCGTGCCAGCTGCATTAAT |
| F_UA1 | TTAATGCAGCTGGCACGACAGCGGCCGCAGCTGGCGGAGATCACCTTG |
| R_UA1 | ACTCCCGCATCTTACCCTGAATATTCAGGGTAAGC |
| F_LA1 | TCAGGGTAAGATGCGGGAGTGGGGGTGA |
| R_LA1 | TCGGGGCTGGCTTAACTATGGCGGCCGCATTCACTGGCATCGCACCG |
| F_SS9gRNA | ATCCTTGACAGCTAGCTCAGTCCTAGGTATAATACTAGTTCTGGCGCAGTTGATATGTA |
| R_SS9gRNA | ACTAGCCTTATTTTAACTTGCTATTTCTAGCTCTAAAACTACATATCAACTGCGCCAGA |
| F_UA2 | TTAATGCAGCTGGCACGACAGAATTCCCGACGTCCATCCAGCCC |
| F_UA2 | GATATAATAAACCTGTTTGACATATCAACTGCGCCAGAGG |
| F_LA2 | TCAAACAGGTTTATTATATCGCGTTGATTATTGATGC |
| R_LA2 | TCGGGGCTGGCTTAACTATGGAATTCATAACCCGCCACAGTAGTTC |
| F_bb3 | TACCAGTGGCGGCTAGTCTTGGACTCCTGTTG |
| R_bb3 | TCGCCGCCTGTGCAGGATTAGCAGATGTGTG |
| F_eut1 | TCTGCTAATCCTGCACAGGCGGCGACTCATG |
| R_eut1 | GCGTAATCGCCCGCTGGGAGACTTTTTTCGCC |
| F_eut2 | AAAGTCTCCCAGCGGGCGATTACGCTGCTCAAC |
| R_eut2 | ATGCGCGCCCGCGCCAATCACCGTGGCGCGC |
| F_eut3 | CACGGTGATTGGCGCGGGCGCGCATACCCT |
| R_eut3 | AGTCCAAGACTAGCCGCCACTGGTACGCTGGG |

**Supplementary Table 1**. Primers used in the present study.

| Position | Protein | Gene orientation | UniProt Id | Annotated Pfam domains | Available information | Present in *S. enterica* LT2? | % Identity | % Similarity |
|---|---|---|---|---|---|---|---|---|
| Upstream *eut* | NudK | Plus | P37128 | PF00293, NUDIX domain | GDP-mannose hydrolase[1]. May play a role in biofilm formation or growth onto solid substrates. | Yes | 88 | 95 |
| | YpfG | Plus | P76559 | PF06674, Protein of unknown function (DUF1176) | Unknown. | Yes, but followed by a gene insertion in the *ypfG-tktB* intergenic region encoding a putative cytoplasmic protein (Uniprot: Q7CQ26) | 73 | 82 |
| | TktB | Minus | P33570 | PF00456, Transketolase thiamine diphosphate binding domain<br>PF02779, Transketolase pyrimidine binding domain<br>PF02780, Transketolase C-terminal domain | Participates in the non-reductive branch of the Pentose Phosphate Pathway[2]. Expression positively regulated by the $\sigma^S$ stress-response sigma factor[3]. | Yes | 93 | 96 |
| | TalA | Minus | P0A867 | PF00923, Transaldolase/Fructose-6-phosphate aldolase | Participates in the non-reductive branch of the Pentose Phosphate Pathway[2]. Expression positively regulated by the $\sigma^S$ stress-response sigma factor[3]. | Yes | 90 | 94 |
| | MaeB | Plus | P76558 | PF00390, Malic enzyme N-terminal domain<br>PF03949, Malic enzyme NAD binding domain<br>PF01515, Phosphate acetyl/butaryl transferase | NADP dependent malic enzyme[4]. Catalyzes the reductive decarboxylation of malate into pyruvate. Ensures a link between the TCA cycle and lower glycolysis. | Yes, but followed by a gene insertion in the *maeB-eutS* intergenic region encoding an IS200 family transposase | 97 | 94 |
| *eut* operon | EutS | Plus | P63746 | PF00936, BMC domain | Hexameric BMC shell protein. | Yes | 95 | 99 |
| | EutP | Plus | P0A208 | PF10662, RAS-like GTPase superfamily domain | Probable acetate kinase.<br>May play a role in positioning the Eut BMCs. | Yes | 84 | 91 |
| | EutQ | Plus | P76555 | PF06249, EutQ | Probable acetate kinase. | Yes | 88 | 93 |
| | EutT | Plus | P65643 | PF01923, Cobalamin adenosyltransferase | Converts cyanocobalamin to adenosylcobalamin. | Yes | 88 | 93 |
| | EutD | Plus | P77218 | PF01515, Phosphate acetyl/butaryl transferase | Phosphate acetyltransferase | Yes | 89 | 95 |
| | EutM | Plus | P41791 | PF00936, BMC domain | Hexameric BMC shell protein. | Yes | 96 | 97 |
| | EutN | Plus | P0AEJ8 | PF03319, EutN/carboxysome | Pentameric BMC shell protein. | Yes | 88 | 93 |
| | EutE | Plus | P77445 | PF00171, Aldehyde dehydrogenase family | Acetaldehyde dehydrogenase. | Yes | 94 | 97 |
| | EutJ | Plus | P77277 | PF14450, Cell division protein FtsA | Chaperone of unknown function. | Yes | 89 | 94 |
| | EutG | Plus | P76553 | PF00465, Iron-containing alcohol dehydrogenase | Alcohol dehydrogenase. | Yes | 81 | 86 |
| | EutH | Plus | P76552 | PF04346, EutH | Ethanolamine transporter. | Yes | 95 | 96 |
| | EutA | Plus | P76551 | PF06277, EutA | Ethanolamine ammonia-lyase reactivase. | Yes | 90 | 95 |
| | EutB | Plus | P0AEJ6 | PF06751, EutB | Ethanolamine-ammonia lyase large subunit | Yes | 98 | 99 |
| | EutC | Plus | P19636 | PF05985, EutC | Ethanolamine-ammonia lyase small subunit | Yes | 91 | 94 |
| | EutL | Plus | P76541 | PF00936, BMC domain | Trimeric BMC shell protein | Yes | 94 | 98 |
| | EutK | Plus | P76540 | PF00936, BMC domain<br>PF16365, EutK C-terminus | BMC shell protein with a potential DNA binding motif | Yes | 78 | 83 |
| | EutR | Plus | P36547 | PF12833, Helix-turn-helix domain | DNA-binding transcriptional activator | Yes, but followed by 2 gene insertions in the *eutR-hemF* intergenic region (Uniprot: Q9ZFU5 and Q9ZFU6) | 92 | 97 |
| Downstream *eut* | HemF | Minus | P36553 | PF01218, Coproporphyrinogen III oxidase | Replaces HemN in the heme biosynthetic pathway when peroxide is present[5]. Only functions under aerobic conditions. | Yes | 92 | 96 |
| | AmiA | Minus | P36548 | PF01520, N-acetylmuramoyl-L-alanine amidase | Plays a role in cell wall peptidoglycan recyling[6]. Involved in the septation process during cell division. | Yes | 88 | 95 |
| | YpeA | Plus | P76539 | PF00583, Acetyltransferase (GNAT) family | Unknown. Does not seem to act as a lysine acetyl transferase[7]. | Yes | 94 | 98 |
| | YfeZ | Plus | P76538 | PF11143, Protein of unknown function (DUF2919) | Unknown. Putative inner membrane protein. | Yes | 74 | 81 |
| | YfeY | Plus | P76537 | PF06572, Protein of unknown function (DUF1131) | Unknown. Expression regulated by the $\sigma^E$ envelope stress-response sigma factor[8]. | Yes | 82 | 91 |
| | YfeX | Plus | P76536 | PF04261, Dyp-type peroxidase N-terminal | Converts protoporphyrinogen IX and coproporphyrinogen III into porphyrins[9]. Expressed under anaerobic conditions. May be required for the synthesis of some components of the anaerobic respiratory chain. | Yes | 93 | 96 |
| | YfeW | Minus | P77619 | PF00144, Beta-lactamase | Penicillin binding protein PBP4B[10]. Unknown function. | Yes, but not at the same genomic location (in the intergenic region upstream *nudK*) | 73 | 84 |
| | MurP | Minus | P77272 | PF00367, Phosphotransferase system EIIB<br>PF02378, Phosphotransferase system EIIC | N-acetylmuramic acid/anhydro-N-acetylmuramic acid transporter. Plays a role in cell wall peptidoglycan recycling[11]. | No | ND | ND |

**Supplementary Table 2.** Proteins encoded within the EUT1 locus of *E. coli* K-12 W3110. Proteins encoded within the *eut* operon are highlighted in orange. Sequence alignments were performed using BlastP to determine the percentages of primary structure identity and similarity with the EUT1 locus proteins of *Salmonella enterica* subsp. *enterica* serovar Typhimurium LT2.

| Protein | Fold-change | p-value | Predicted function |
|---|---|---|---|
| eutP | 368,9 | 1,22E-09 | Probable acetate kinase |
| eutM | 325,1 | 8,87E-08 | Hexameric BMC shell protein |
| eutT | 206,6 | 6,23E-09 | Corrinoid adenosyltransferase |
| eutE | 160,9 | 1,41E-07 | Acetaldehyde dehydrogenase |
| amtB | 150,6 | 3,42E-07 | Ammonium transporter, AMT family |
| eutS | 148,6 | 1,35E-07 | Hexameric BMC shell protein |
| eutQ | 134,1 | 9,01E-09 | Probable acetate kinase |
| eutB | 110,5 | 3,13E-09 | Ethanolamine-ammonia lyase large subunit |
| eutG | 84,8 | 1,18E-08 | Alcohol dehydrogenase |
| glnK | 73,5 | 1,47E-06 | Nitrogen assimilation regulatory protein for GlnL, GlnE and AmtB |
| eutC | 66,1 | 8,99E-09 | Ethanolamine-ammonia lyase small subunit |
| eutH | 65,7 | 4,73E-08 | Ethanolamine permease |
| eutD | 58,9 | 7,28E-08 | Phosphate acetyltransferase |
| eutL | 48,3 | 9,27E-08 | Trimeric BMC shell protein |
| eutR | 29,8 | 3,70E-05 | DNA-binding transcriptional activator |
| eutK | 22,6 | 3,69E-08 | BMC shell protein |
| eutA | 19,6 | 1,22E-07 | Ethanolamine ammonia-lyase reactivase |
| eutN | 15,5 | 5,74E-07 | Pentameric BMC shell protein |
| priA | 7,1 | 1,80E-06 | Primosomal protein n' (replication factor y) |
| ddpX | 7,0 | 3,74E-03 | D-Ala-D-Ala dipeptidase, Zn-dependent |
| acrB | 6,8 | 4,01E-07 | Multidrug efflux pumpRND permease |
| hrpA | 6,1 | 9,99E-06 | ATP-dependent RNA helicase |
| ddpA | 6,0 | 1,79E-06 | Putative d,d-dipeptide abc transporter periplasmic binding protein |
| eutJ | 5,4 | 8,05E-08 | Putative chaperone |
| oppA | 5,1 | 7,28E-05 | Oligopeptide abc transporter periplasmic binding protein |
| uxuA | 4,3 | 1,06E-06 | D-mannonate dehydratase |
| guaD | 4,1 | 1,40E-06 | Guanine deaminase |
| ampC | 3,9 | 3,17E-06 | β-lactamase |
| cbl | 3,7 | 3,81E-04 | DNA-binding transcriptional activator Cbl |
| ybiO | 3,7 | 1,18E-04 | Mechanosensitive channel |
| gcvP | 3,5 | 4,33E-10 | Glycine decarboxylase |
| glnL | 2,9 | 5,38E-06 | Two-component system, ntrc family |
| yqjH | 2,9 | 1,80E-02 | NADPH-dependent ferric chelate reductase |
| cycA | 2,8 | 9,46E-06 | D-serine/alanine/glycine/:$H^+$symporter |
| rsmA | 2,7 | 1,21E-07 | rRNA dimethyltransferase |
| astD | 2,6 | 1,35E-07 | Succinylglutamate-semialdehyde dehydrogenase |
| glnA | 2,6 | 2,13E-06 | Glutamine synthetase |
| astC | 2,6 | 1,53E-05 | Succinylornithine transaminase |
| glnG | 2,6 | 4,28E-06 | Two-component system, ntrc family |
| rnpA | 2,4 | 4,14E-06 | Ribonuclease P protein |
| asnB | 2,4 | 1,85E-06 | Asparagine synthase (glutamine-hydrolysing) |
| hisQ | 2,3 | 1,65E-03 | Lysine/arginine/ornithine ABC transporter, membrane subunit |
| gcvH | 2,3 | 4,68E-02 | Glycine cleavage system H protein |
| argT | 2,3 | 1,48E-06 | Lysine/arginine/ornithine ABC transporter, periplasmic binding protein |
| ygiQ | 2,2 | 1,26E-04 | Protein of unknown function |
| dgcM | 2,1 | 3,51E-05 | Diguanylate cyclase |
| asnA | 2,1 | 3,43E-04 | Aspartate-ammonia ligase |
| fbp | 2,0 | 2,96E-04 | Fructose-1,6-bisphosphatase class 1 |
| yciF | 0,5 | 2,28E-05 | Putative rubrerythrin/ferritin-like metal-binding protein |
| glaH | 0,4 | 1,63E-03 | Glutarate dioxygenase |
| ybjP | 0,4 | 4,96E-02 | DUF3828 domain-containing lipoprotein |
| metF | 0,4 | 1,26E-05 | Methylenetetrahydrofolate reductase |
| phoA | 0,4 | 5,52E-05 | Alkaline phosphatase |
| lldD | 0,2 | 1,80E-06 | L-lactate dehydrogenase |
| metE | 0,1 | 1,89E-10 | Cobalamin-independent homocysteine transmethylase |
| nhaA | 0,0 | 1,10E-02 | $Na^+$:$H^+$ antiporter |

**Supplementary Table 3.** List of the *E. coli* K12 W3110 WT proteins that were differentially accumulated in M9 glycerol EA B12 compared to M9 glycerol $NH_4Cl$ (fold-change > 2 or < 0,5 and p-value < 0,05). The annotated functions as indicated on STRING v11.5 are also reported.

| Protein | Fold-change | p-value | Predicted function |
|---|---|---|---|
| NudK | ND | ND | GDP-mannose hydrolase |
| YpfG | ND | ND | Unknown |
| TktB | 0,69 | 4,27E-05 | Transketolase |
| TalA | 0,68 | 1,77E-04 | Transaldolase |
| MaeB | 1,00 | 8,78E-01 | Malic enzyme |
| HemF | ND | ND | Coproporphyrinogen III oxidase |
| AmiA | 1,20 | 2,63E-02 | N-acetylmuramoyl-L-alanine amidase |
| YpeA | 0,96 | 1,29E-01 | Acetyltransferase (GNAT) family |
| YfeZ | ND | ND | Unknown |
| YfeY | 0,87 | 1,09E-02 | Unknown |
| YfeX | 0,88 | 2,05E-02 | Dye-decolorizing peroxidase |

**Supplementary Table 4.** Abundance of the detected EUT1 locus ancillary proteins in M9 glycerol EA B12 compared to M9 glycerol $NH_4Cl$. None of the detected EUT1 locus ancillary proteins passed the set significance thresholds (fold-change > 2 or < 0,5 and p-value < 0,05). ND: not detected.

| Reaction | Description | Localization | Value |
|---|---|---|---|
| **Experimentally determined values** | | | |
| μ | Growth rate ($h^{-1}$) | NA | 0.37 ± 0.01 |
| $q_{S)Glycerol}$ | Glycerol uptake from extracellular medium | Extracellular | 13.68 ± 0.15 |
| $q_{S)Ethanolamine}$ | Ethanolamine uptake from extracellular medium | Extracellular | 7.04 ± 0.23 |
| $q_{P)Ethanol}$ | Ethanol excretion into extracellular medium | Cytosol | 2.67 ± 0.11 |
| $q_{P)Acetate}$ | Acetate excretion into extracellular medium | Cytosol | 1.64 ± 0.09 |
| $q_{P)NH4}$ | Ammonium excretion into extracellular medium | Extracellular | 0.86 ± 0.21 |
| **Predicted values for ethanolamine-derived acetaldehyde metabolism** | | | |
| $v_{EutBC}$ | EAL | BMC | 7.04 ± 0.23 |
| $v_{eBMC)Acetaldehyde}$ | Acetaldehyde leakage from the Eut BMCs | BMC | 2.67 ± 0.11 |
| $v_{EutG}$ | Acetaldehyde to ethanol conversion ($NAD^+$-dependent) | BMC | 2.67 ± 0.11 |
| $v_{eBMC)Ethanol}$ | Ethanol diffusion out of the BMC | BMC | 2.67 ± 0.11 |
| $v_{EutE}$ | Acetaldehyde to acetyl-CoA conversion (NADH-dependent) | BMC | 2.67 ± 0.11 |
| $v_{EutD}$ | Acetyl-CoA to acetyl-P conversion | BMC | 2.67 ± 0.11 |
| $v_{eBMC)Acetyl-P}$ | Acetyl-P diffusion out of the BMC | BMC | 2.67 ± 0.11 |
| **Predicted values for glycerol metabolism** | | | |
| $V_{gly)biomass}$ | Biomass production from glycerol | Cytosol | 11.17 ± 0.20 |
| $V_{gly)glycolysis}$ | Acetyl-P production from glycerol | Cytosol | 2.51 ± 0.06 |
| **Predicted values for cytosolic acetyl-P metabolism** | | | |
| $v_{EutP, EutQ, AckA}$ | Acetyl-P to acetate conversion | Cytosol | 1.64 ± 0.09 |
| $v_{Pta}$ | Acetyl-P to acetyl-CoA (to biomass) conversion | Cytosol | 3.54 ± 0.14 |
| **Predicted values for ethanolamine-derived ammonium metabolism** | | | |
| $v_{eBMC)NH4}$ | Ammonium diffusion out of the BMC | BMC | 7.04 ± 0.23 |
| $v_{NH4)biomass}$ | Biomass production from ammonium | Cytosol | 6.18 ± 0.08 |

**Supplementary Table 5.** Growth parameters of *E. coli* K12 W3110 WT cultivated aerobically in M9 medium containing $^{12}C_2$-glycerol, $^{13}C_2$-EA and vit B12. The experimentally determined substrate uptake ($q_S$) and product excretion ($q_P$) rates are expressed in mmol.(gDW.h)$^{-1}$. Intracellular flux values (v) determined using the isotopic model are also expressed in mmol.(gDW.h)$^{-1}$. DW: dry weight; NA: not appliable. Data is the average of n=3 independent replicates, errors indicate SD.

### Supplementary figures

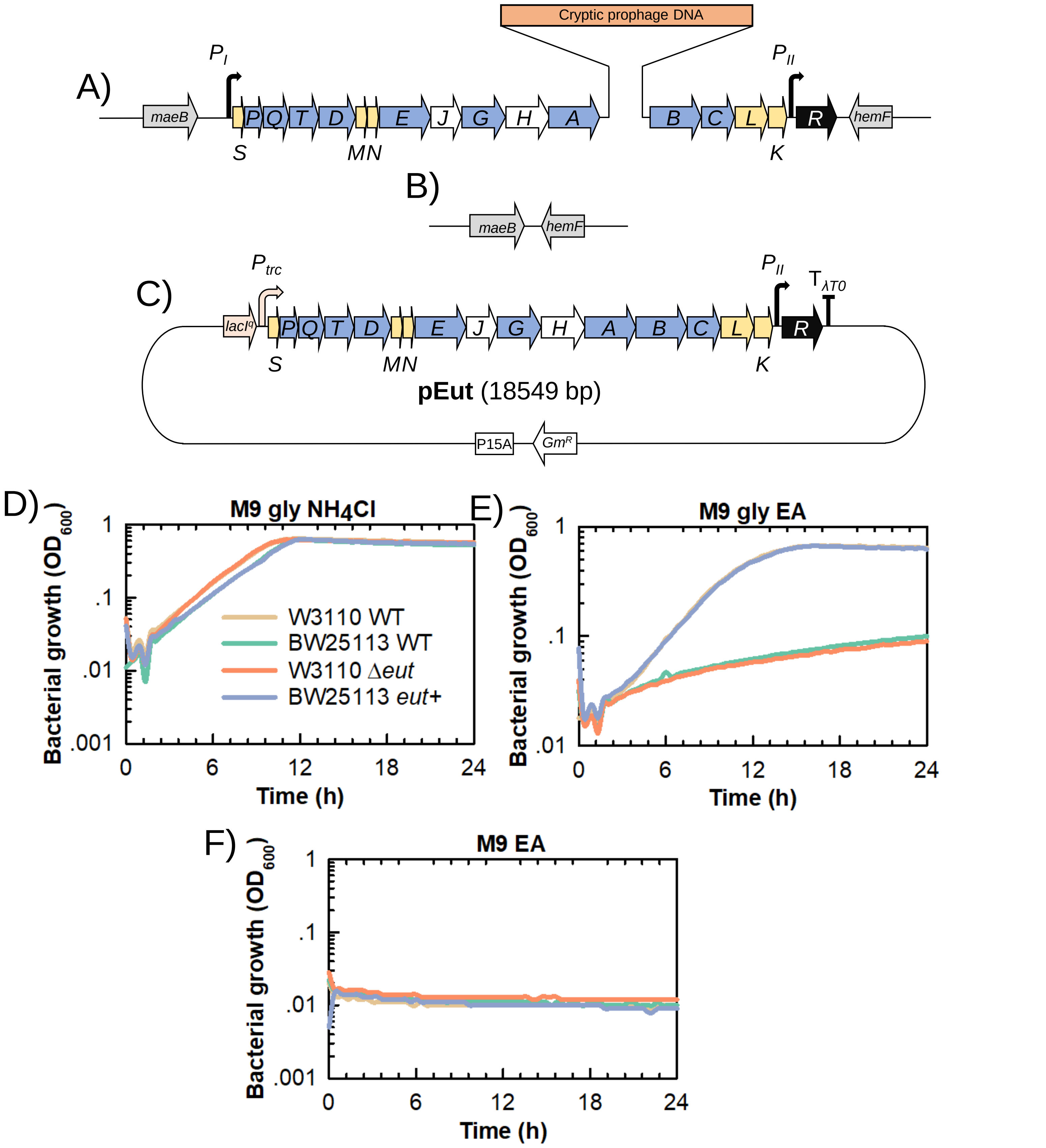


**Figure S1. Capacity to utilize EA as a N source in *Escherichia coli* K12 and mutant derivatives.** Schematic representation of the *eut* operon in A) the chromosome of BW25113 WT where it harbors a 6,9 kb prophage DNA insertion downstream *eutA*, B) the chromosome of W3110 *Δeut* where the entire operon was scarlessly removed by genome editing and C) the pEut plasmid where the *eut* operon from W3110 WT was cloned into pSEVA661 under control of the IPTG-inducible $P_{trc}$ promoter. The BW25113 *eut*+ mutant was generated by scarlessly removing the prophage DNA insertion through genome editing, thus reconstituting an uninterrupted *eut* operon (yielding the same *eut* operon as in W3110 WT). D) to F) Plate reader measurement of optical density at 600 nm over time. The strains were grown in D) M9 derivative with glycerol as the C source and $NH_4Cl$ as the N source; E) M9 derivative with glycerol as the C source and EA as the sole N source, including also vitamin B12; F) M9 derivative with EA (160 mM) as the C and N source, including also 200 nM vitamin B12. Data shows a representative experiment out of at least n = 3 independent replicates.

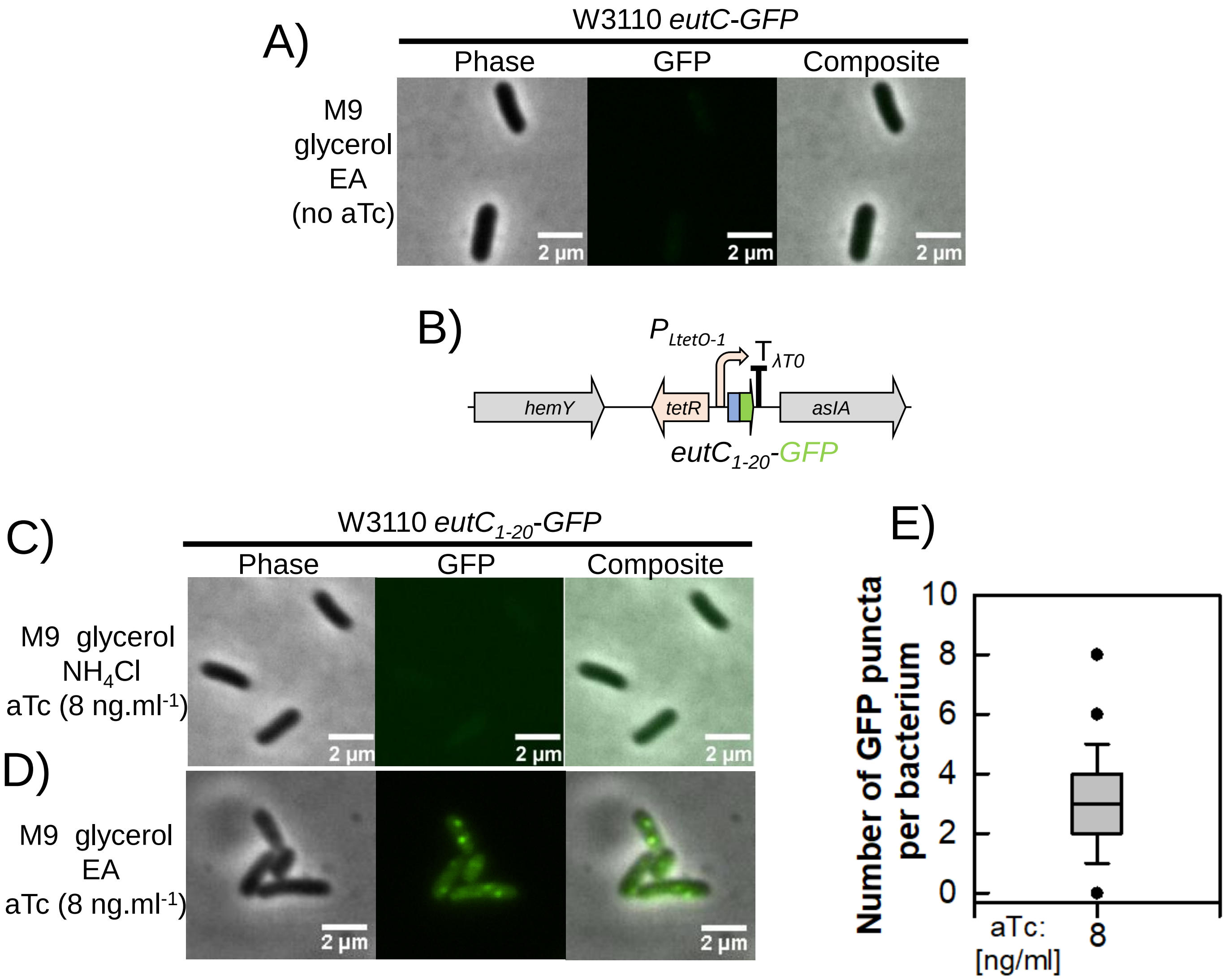


**Figure S2. Epifluorescence microscopy to determine the EutC-GFP and EutC$_{1\text{-}20}$-GFP subcellular localization.** A) Representative micrograph showing a phase contrast image, GFP fluorescence signal as well as a composite image from W3110 *eutC-GFP* grown in M9 glycerol EA vit B12 without any aTc added. B) Schematic representation of the P$_{LtetO\text{-}1}$::eutC$_{1\text{-}20}$-GFP expression cassette. This construct was inserted at the SS9 safe locus site in W3110 WT to yield W3110 *eutC$_{1\text{-}20}$-GFP*. C) and D) Representative micrographs showing phase contrast images, GFP fluorescence signal as well as composite images from W3110 *eutC$_{1\text{-}20}$-GFP* grown in C) M9 glycerol $NH_4Cl$ with 8 ng.ml$^{-1}$ aTc and D) M9 glycerol EA vit B12 with 8 ng.ml$^{-1}$ aTc. E) Box plot indicating the number of GFP puncta per bacterium out of n = 30 individual cells for cultures grown in M9 glycerol EA vit B12 with 8 ng.ml$^{-1}$ aTc.

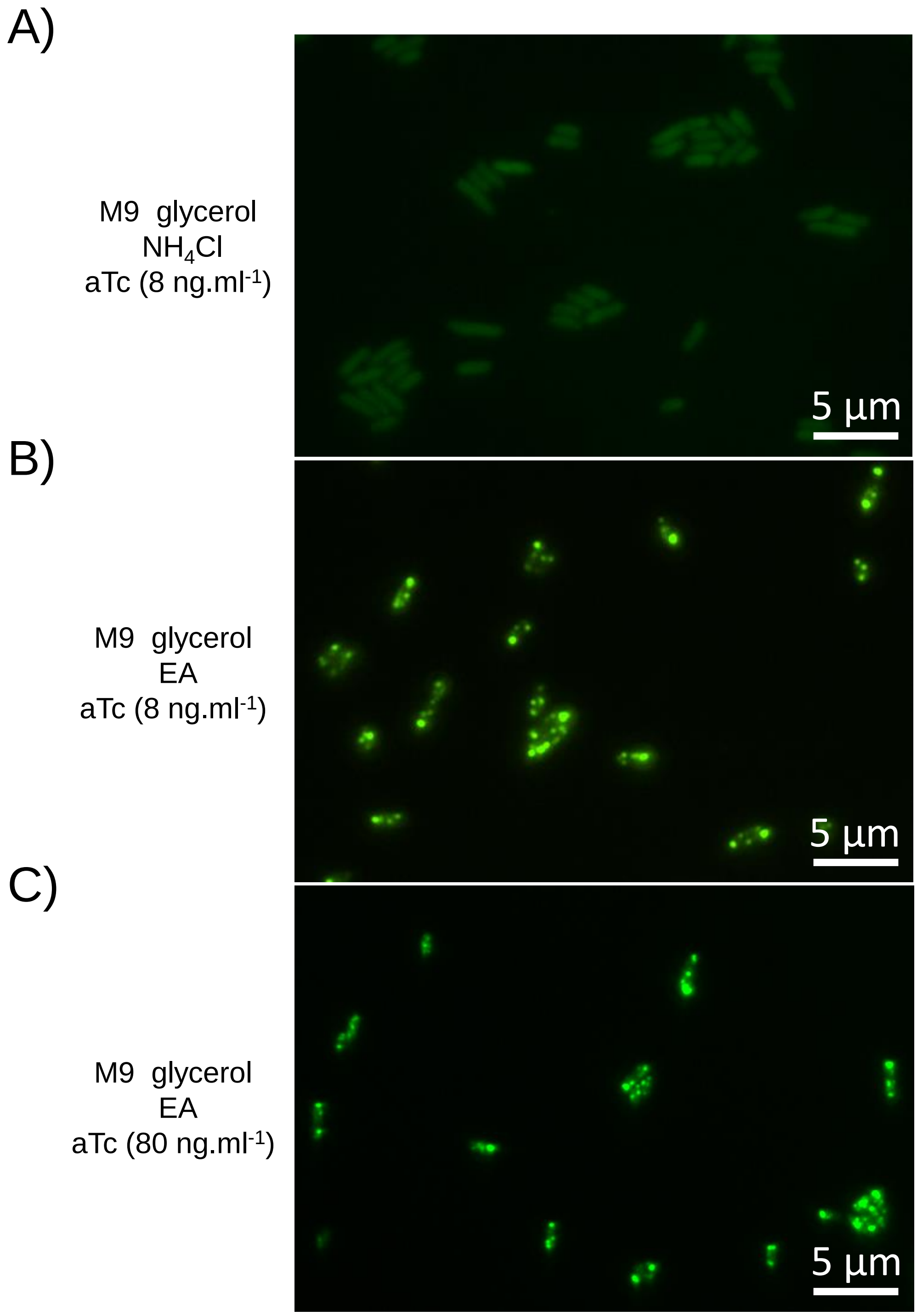


**Figure S3. Wider views of the GFP channel for W3110 eutC-GFP grown in various media.** Micrographs showing the strain grown in A) M9 glycerol $NH_4Cl$ with 8 $ng.ml^{-1}$ aTc, B) M9 glycerol EA vitamin B12 with 8 $ng.ml^{-1}$ aTc and C) M9 glycerol EA vitamin B12 with 80 $ng.ml^{-1}$ aTc. Scale bars indicate 5 µm.

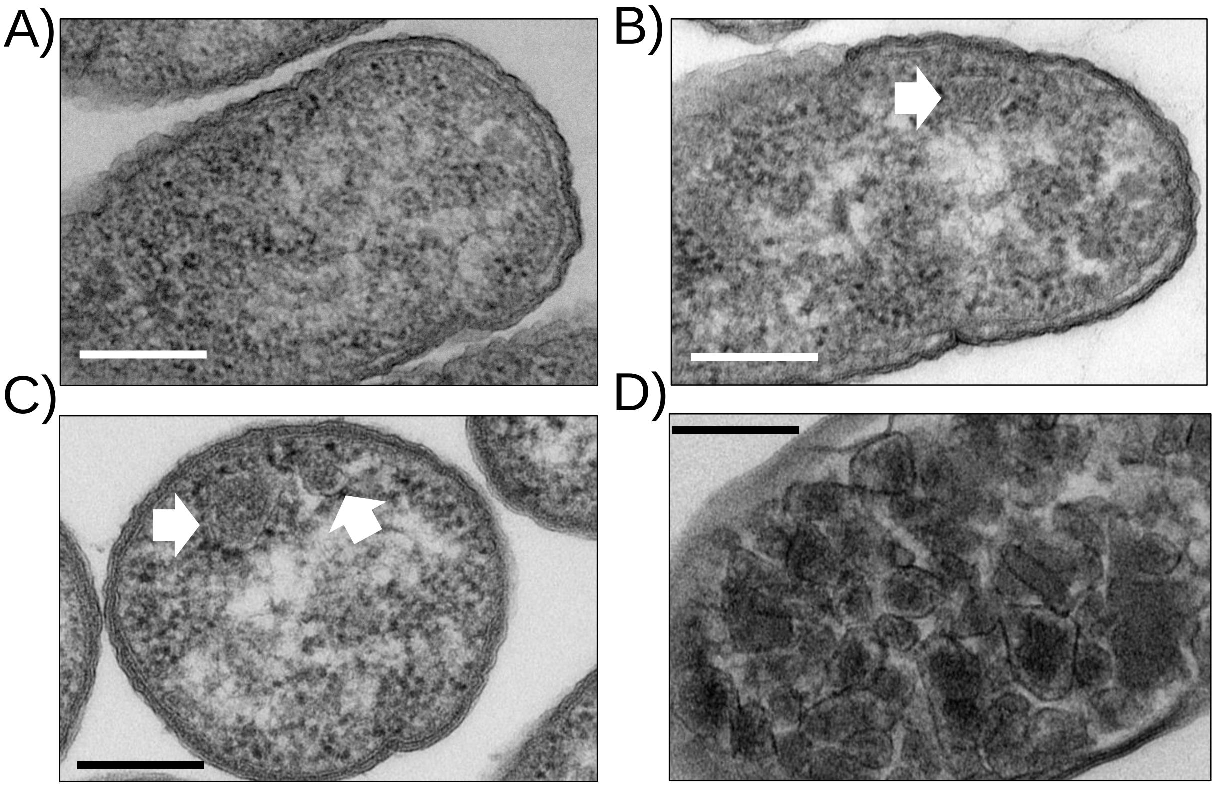


**Figure S4. Observation of the *Escherichia coli* K12 W3110 Eut BMCs by TEM.** A) W3110 WT in M9 glycerol $NH_4Cl$; B) W3110 WT in M9 glycerol EA vit B12; C) W3110 WT in LB EA vit B12 and D) W3110 Δeut pEut_WT in LB 20 µM IPTG. The filled white arrows indicate Eut BMCs in B and C. Scale bars show 200 nm.

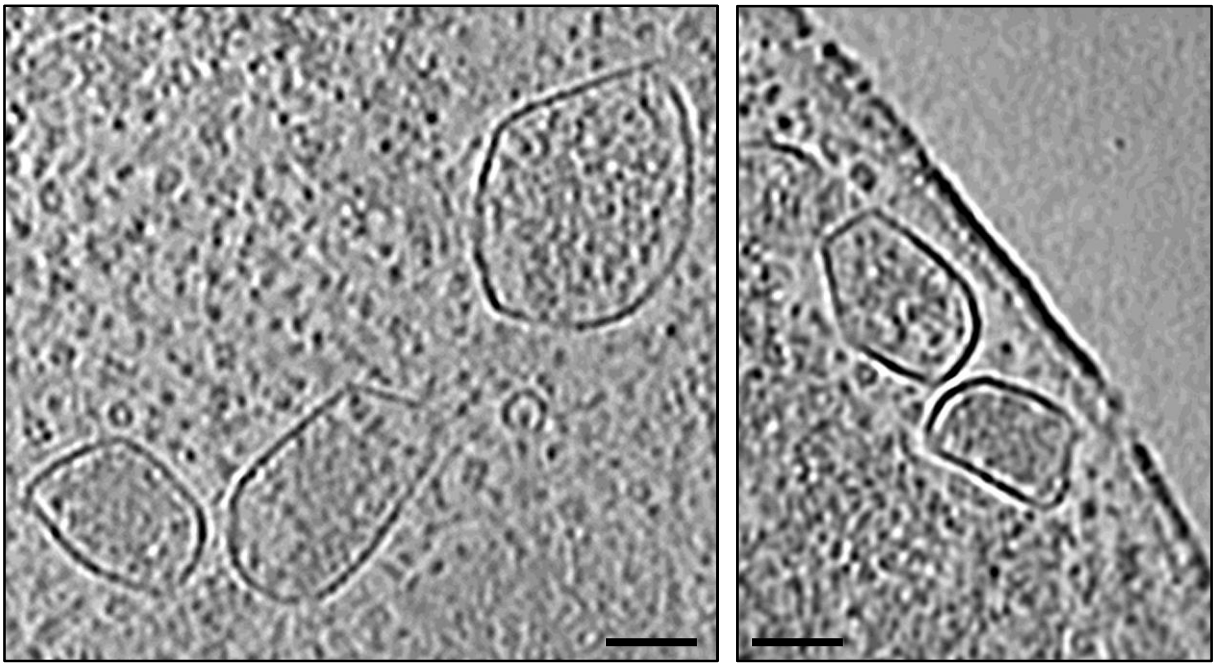

**Figure S5. CryoET observation of the Eut BMCs formed by W3110 Δ*eut* pEut.** The *eut* overexpressor strain was grown in M9 glycerol EA with 20 µM IPTG. Representative electron cryotomogram slices are shown. The scale bars indicate 50 nm.

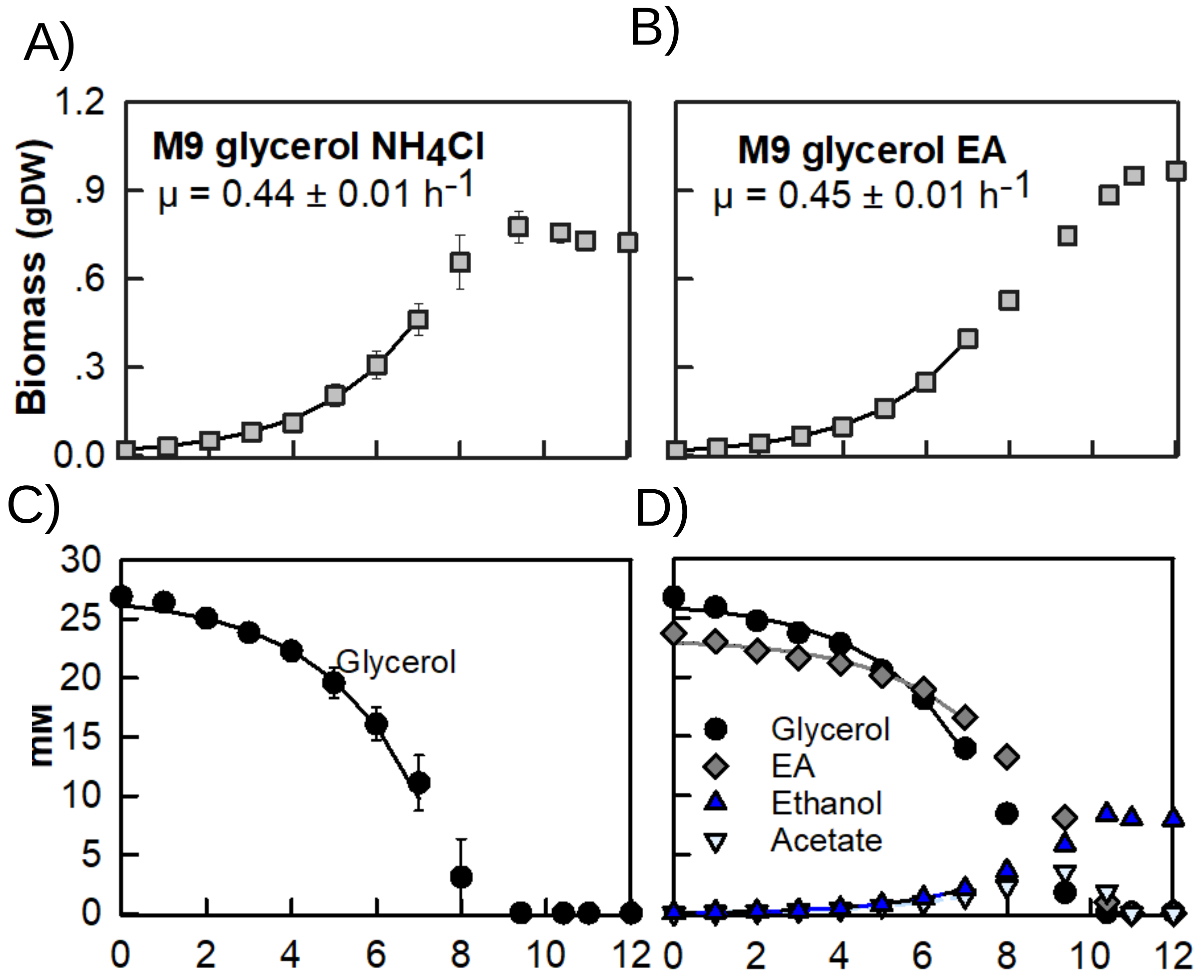


**Figure S6. Physiological characterization of *Escherichia coli* K12 W3110 WT grown in M9 glycerol containing $NH_4Cl$ or EA as the sole N source.** Cultures were grown aerobically in A) and C) M9 glycerol (30 mM) $NH_4Cl$ (25 mM) or B) and D) M9 glycerol (30 mM) EA (25 mM) vit B12 (200 nM). A) and B) Biomass accumulation (grams dry weight, gDW); C) and D) Exometabolome profiling by $^{1}H$-NMR. Data is the average of n = 3 biological replicates, bars show SD. The solid lines are the best fits obtained with PhysioFit.

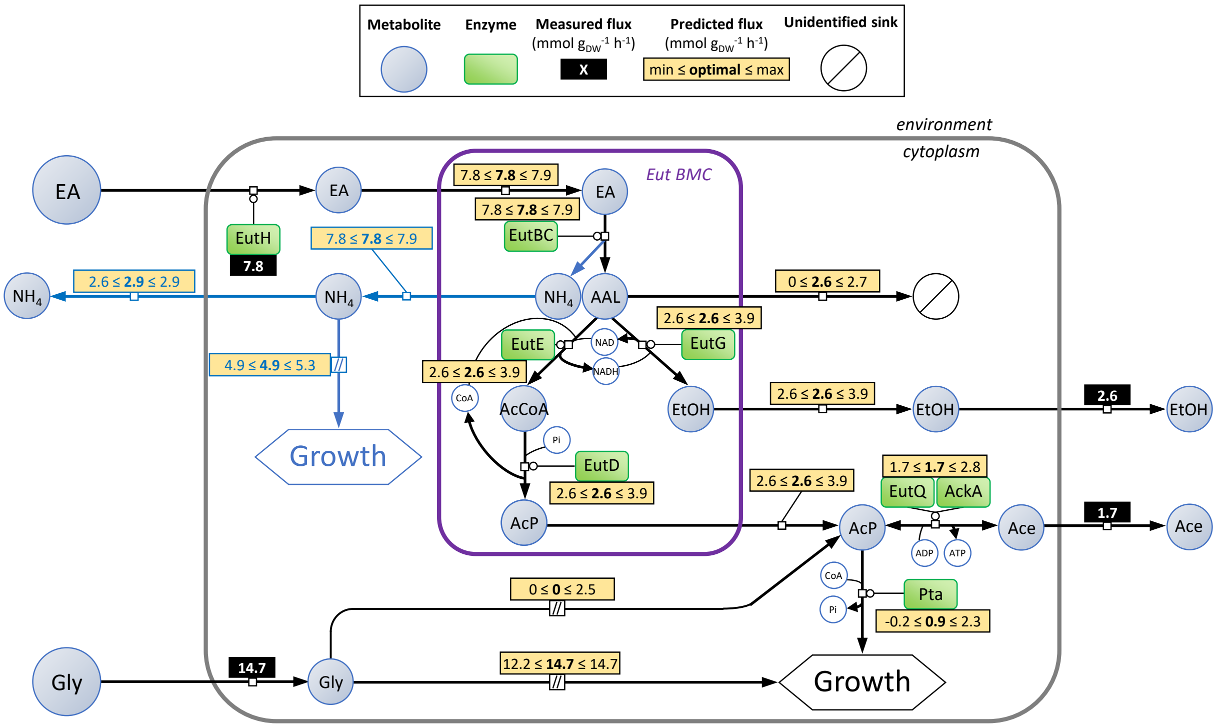


**Figure S7. FBA/FVA analysis of EA metabolism in *E. coli* K-12 W3110.** Black rectangles indicate the experimentally determined exchange fluxes for W3110 WT grown in M9 glycerol (30 mM) EA (25 mM) (data from Fig.S5). Yellow rectangles report the predicted fluxes. The BMC-associated fluxes were deduced assuming that NADH and CoA-SH are recycled internally while ATP is recycled within the cytosol. We also assumed that acetate and ethanol were produced from the BMCs. The non-constrained fluxes were simulated by flux balance analysis (optimal value: bold font) combined with flux variability analysis (to determine possible value ranges while maintaining 99% of the fitness). Our model aimed at maximizing ATP production while the growth rate was set to 0.45 $h^{-1}$ to match the experimental data. Flux values are given in $mmol.g_{DW}^{-1}.h^{-1}$. AAL: acetaldehyde; AcCoA: acetyl-CoA; AcP: acetyl-phosphate; EtOH: ethanol; Gly: glycerol; $NH_4$: ammonium.

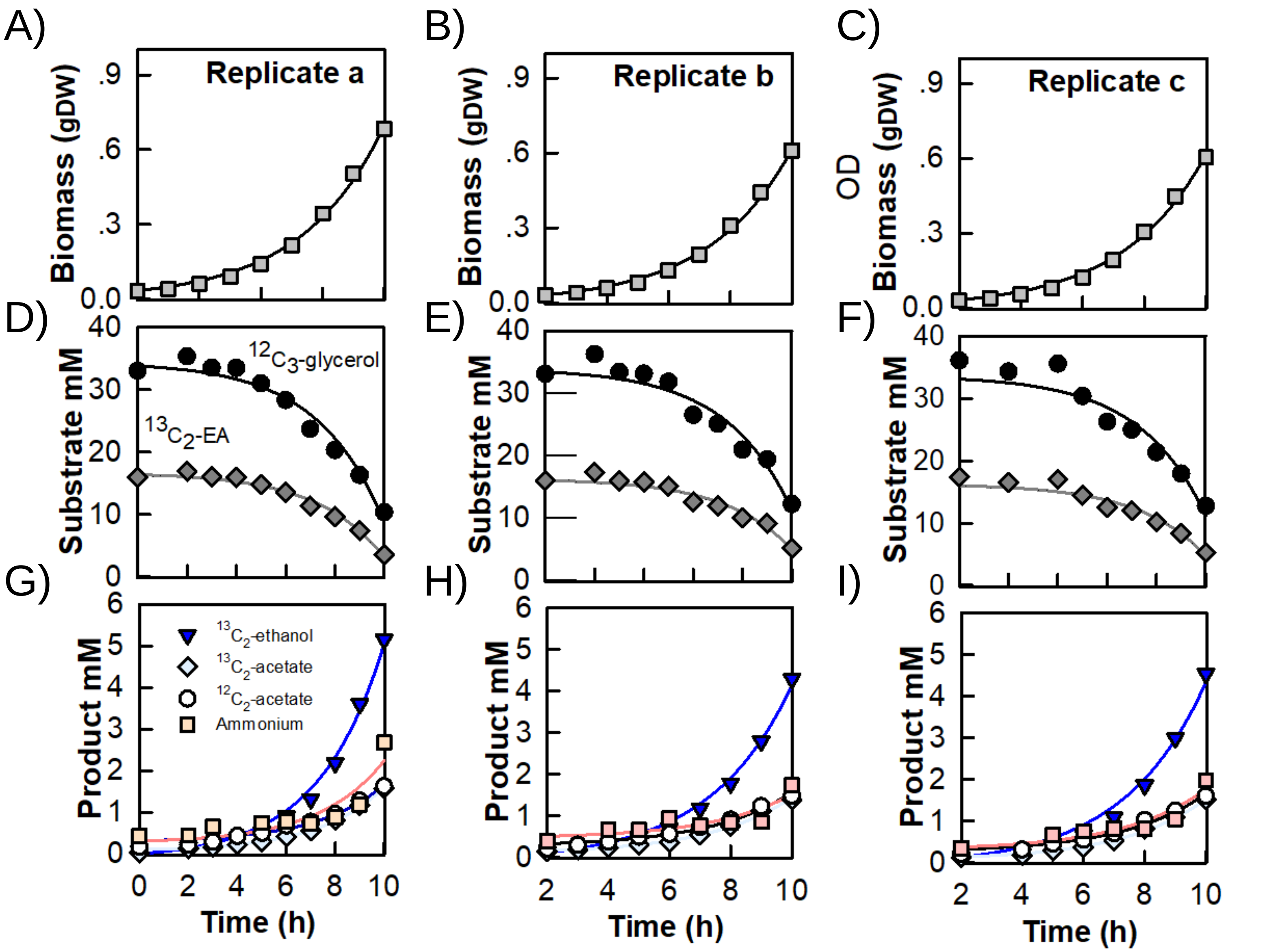


**Figure S8. The dynamic isotopic model fits the experimental data satisfactorily.** W3110 WT cultures (3 biological replicates named a, b and c) were grown aerobically in M9 medium containing $^{12}C_3$-glycerol and $^{13}C_2$-EA. A), B) and C) Biomass accumulation (grams dry weight, gDW) in biological replicates a, b and c. Exometabolome composition was determined by $^{1}$H-NMR. D), E) and F) $^{12}C_3$-glycerol and $^{13}C_2$-EA uptake in biological replicates a, b and c. G), H) and I) $^{13}C_2$-EtOH, $^{13}C_2$-acetate, $^{12}C_2$-acetate and ammonium production in biological replicates a, b and c. Solid lines show the best fits with our dynamic isotopic model. The experimental data from all 3 replicates were averaged to produce Figure 5.

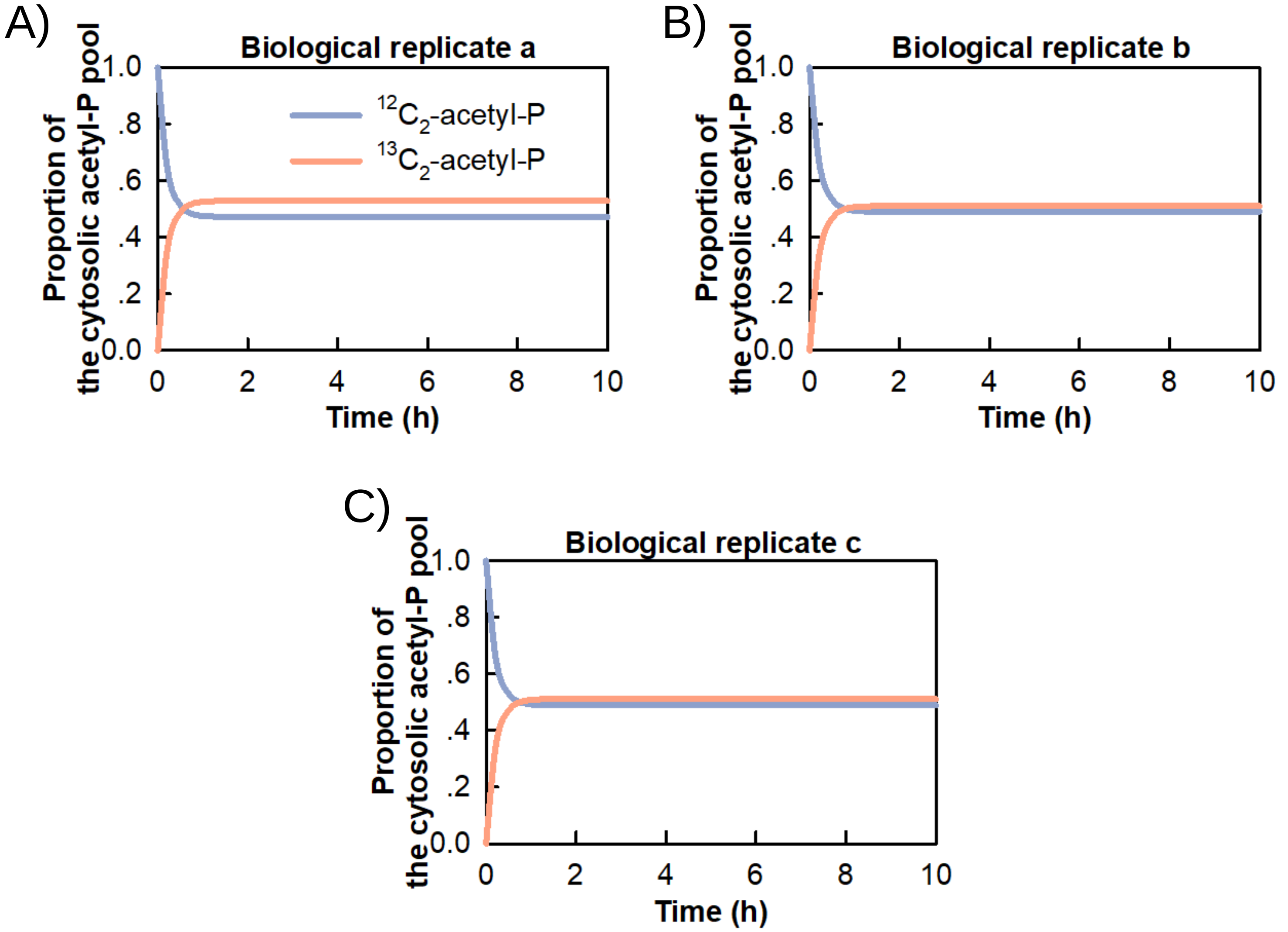


**Figure S9. Predicted evolution of the cytosolic acetyl-P pool isotopic composition.** The isotopic model was used to fit data from Fig.5. The predicted proportion of cytosolic $^{12}C_2$-acetyl-P (derived from glycerol) VS $^{13}C_2$-acetatyl-P (derived from EA) over time is shown for each biological replicate (A to C).
