## Supplementary Data 2 for "Integrative *in vivo* analysis of the ethanolamine utilization bacterial microcompartment in *Escherichia coli.*"

### Isotopic model of ethanolamine and glycerol co-metabolism in *Escherichia coli*

-

Documentation

### 1. Model overview

This model contains a coarse-grained representation of the central metabolism and of the Eut bacterial microcompartments (BMCs) of *Escherichia coli*. We used this model to investigate ethanolamine and glycerol co-metabolism in *E. coli*.

This model was developed with COPASI (http://copasi.org). All models are available in SBML and COPASI formats at https://github.com/MetaSys-LISBP/ethanolamine_metabolism. The model can also be downloaded from the Biomodels database (http://www.ebi.ac.uk/biomodels) with identifier MODEL2403010002.

This model comprises 3 compartments (the environment and the cell, with the cytosol and the Eut BMCs), 20 species (19 metabolites and biomass) and 21 reactions that represent the following processes (Figure 1):

- glycerol uptake and glycolytic conversion into acetyl-phosphate
- ethanolamine assimilation and conversion through the eut BMCs
- ammonia, ethanol and acetate excretion
- ethanol evaporation
- anabolic utilization of acetyl-phosphate
- exchange of metabolites between the environment, the cytosol and the eut BMCs
- growth

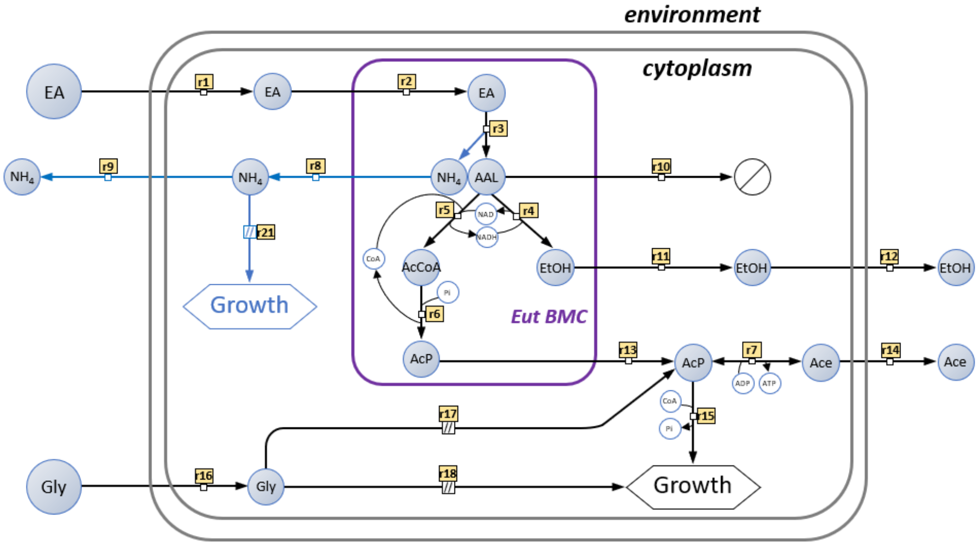


**Figure 1.** Metabolic network of glycerol and ethanolamine co-metabolism in *E. coli*. The diagram follows the conventions of the Systems Biology Graphical Notation process description.

#### 2. Model units

Model units are millimole (mmol) for amounts of metabolites, litre (L) for volumes, and hour (h) for time. Amount of biomass is expressed as gram dry weight ($g_{DW}$).

#### 3. Reactions

The reactions included in the model are listed in the table below.

| ***Name*** | ***Reaction*** | ***Flux*** |
|---|---|---|
| r1_EA_in | EA_e -> EA_c | v1 |
| r2_EA_xch | EA_c -> EA_eut | v2 |
| r3_EutBC | EA_eut -> NH4_eut + AAL_eut | v3 |
| r4_EutG | AAL_eut + NADH_eut -> EtOH_eut + NAD_eut | v4 |
| r5_EutE | AAL_eut + NAD_eut -> AcCoA_eut + NADH_eut | v5 |
| r6_EutD | AcCoA_eut -> AcP_eut | v6 |
| r7_EutQ | AcP_c -> Ace_c | v7 |
| r8_NH4_xch | NH4_eut -> NH4_c | v8 |
| r9_NH4_out | NH4_c -> NH4_e | v9 |
| r10_AAL_sink | AAL_eut -> Ø | v10 |
| r11_EtOH_xch | EtOH_eut -> EtOH_c | v11 |
| r12_EtOH_out | EtOH_c -> EtOH_e | v12 |
| r13_AcP_xch | AcP_eut -> AcP_c | v13 |
| r16_GLY_in | GLY_e -> GLY_c | v14 |
| r17_glycolysis | GLY_c -> AcP_c | v15 |
| r18_GLY_sink | GLY_c -> Ø | v16 |
| r15_AcP_sink | AcP_c -> Ø | v17 |
| r14_Ace_out | Ace_c -> Ace_e | v18 |
| r19_growth | BIOMASS -> 2 * BIOMASS | v19 |
| r20_EtOH_evap | EtOH_e -> Ø | v20 |
| r21_NH4_sink | NH4_c -> Ø | v21 |

#### 4. ODEs system

The differential equations, which describe the progression of the variables over time as a function of the system's rates, balance the concentrations of extracellular species (biomass, glycerol, ethanolamine, ethanol, acetate and ammonia) and of intracellular species (ethanolamine, glycerol, acetaldehyde, ammonia, ethanol, acetate, acetylcoA, NAD(H), acetyl-phosphate).

Extracellular species:

$$\frac{dEA_e}{dt} = -v_1 \cdot X$$

$$\frac{dGLY_e}{dt} = -v_{14} \cdot X$$

$$\frac{dACE_e}{dt} = v_{18} \cdot X$$

$$\frac{dEtOH_e}{dt} = v_{12} \cdot X - v_{20}$$

$$\frac{dNH_{4_e}}{dt} = v_9 \cdot X$$

$$\frac{dX}{dt} = v_{19} \cdot X$$

Intracellular species:

$$\frac{dEA_c}{dt} = v_1 - v_2$$

$$\frac{dEA_{eut}}{dt} = v_2 - v_3$$

$$\frac{dAAL_{eut}}{dt} = v_3 - v_4 - v_5 - v_{10}$$

$$\frac{dNH_{4_{eut}}}{dt} = v_3 - v_8$$

$$\frac{dEtOH_{eut}}{dt} = v_4 - v_{11}$$

$$\frac{dAcCoA_{eut}}{dt} = v_5 - v_6$$

$$\frac{dNADH_{eut}}{dt} = v_5 - v_4$$

$$\frac{dNAD_{eut}}{dt} = v_4 - v_5$$

$$\frac{dAcP_{eut}}{dt} = v_6 - v_{13}$$

$$\frac{dNH_{4_c}}{dt} = v_8 - v_9 - v_{21}$$

$$\frac{dEtOH_c}{dt} = v_{11} - v_{12}$$

$$\frac{dAcP_c}{dt} = v_{13} - v_{14} - v_{15}$$

$$\frac{dGLY_c}{dt} = v_{16} - v_{17} - v_{18}$$

$$\frac{dACE_c}{dt} = v_7 - v_{14}$$

#### 5. Reaction rates

Ethanol evaporation was modelled using a first-order degradation rate law ($v_{20} = k_{evap} \cdot [EtOH]$), as observed experimentally. The rates of all other reactions were defined as constant and constrained to match the steady-state conditions (exponential growth) by balancing the concentration of all intracellular species, resulting in the following equations:

$$v_1 = v_3$$
$$v_2 = v_3$$
$$v_3 = v_4 + v_5 + v_{10}$$
$$v_4 = v_5$$
$$v_5 = v_6$$
$$v_6 = v_7$$
$$v_7 = v_{13}$$
$$v_8 = v_3$$
$$v_9 = v_8 - v_{21}$$
$$v_{11} = v_4$$
$$v_{12} = v_4$$
$$v_{13} = v_{14} + v_{15} - v_{17}$$
$$v_{16} = v_{17} + v_{18}$$

which can be determined from the following fluxes (defined as free parameters): $v_{10}, v_{14}, v_{15}, v_{17}, v_{18}$.

#### 6. Extension with isotopic equations

This dynamic model was extended with isotopic equations as detailed previously (doi: 10.1186/s12918-015-0213-8). Briefly, all reactions (except biomass synthesis) were considered separately for unlabeled and labeled metabolites. For instance, the isotopically extended balance of AcP corresponds to:

$$\frac{dAcP_0}{dt} = v_{17} - \frac{AcP_0}{AcP_0 + AcP_1} \cdot (v_{14} + v_{15})$$
$$\frac{dAcP_1}{dt} = v_{13} - \frac{AcP_1}{AcP_0 + AcP_1} \cdot (v_{14} + v_{15})$$

where subscripts 0 and 1 refers to the unlabeled and labeled metabolites, respectively.

#### 7. Flux calculation

Initial concentrations of unlabelled intracellular metabolites were set to unity, and initial concentrations of labelled metabolites were set to zero. The evaporation rate constant of ethanol was set to its experimental value ($k_{evap} = 0.0379\ h^{-1}$).

Therefore, this model contains 13 free parameters:

- initial concentrations of extracellular glycerol (*GLY*), ethanolamine (*EA*), biomass (*X*), ammonia (*NH₄*), and unlabeled and labeled acetate (*ACE₀* and *ACE₁*)
- acetate production ($v_{14}$), acetyl-phosphate utilization by the rest of metabolism ($v_{15}$), glycolytic conversion of glycerol ($v_{17}$), utilization of a part of glycerol by the rest of metabolism ($v_{18}$), acetaldehyde leak from the BMCs ($v_{10}$), ammonia excretion rate ($v_{21}$) and growth rate ($v_{19}$)

These parameters were estimated by fitting experimental data (time course concentrations of biomass, glycerol, ethanolamine, ammonia, ethanol and of labelled and unlabeled acetate), as detailed in the publication. Values and standard deviations obtained for each of the three independent biological replicates are provided below. All parameters were determined with a good precision in each of the three independent biological replicates. We recalculated all fluxes from the estimated parameters.

| ***Parameter*** | *Replicate 1* | | *Replicate 2* | | *Replicate 3* | | ***mean***[c] | ***sd***[c] |
|---|---|---|---|---|---|---|---|---|
| | *value*[a] | *sd*[b] | *value*[a] | *sd*[b] | *value*[a] | *sd*[b] | | |
| μ | 0.37 | 0.01 | 0.37 | 0.01 | 0.37 | 0.01 | **0.37** | **0.01** |
| $v_{10}$ | 1.72 | 0.43 | 1.73 | 0.53 | 1.62 | 0.64 | **1.69** | **0.06** |
| $v_{14}$ | 1.57 | 0.26 | 1.60 | 0.31 | 1.74 | 0.39 | **1.64** | **0.09** |
| $v_{15}$ | 3.69 | 0.87 | 3.39 | 1.03 | 3.48 | 1.24 | **3.52** | **0.15** |
| $v_{17}$ | 2.63 | 0.86 | 2.43 | 0.96 | 2.55 | 1.15 | **2.54** | **0.10** |
| $v_{18}$ | 11.29 | 0.92 | 11.27 | 1.06 | 10.98 | 1.28 | **11.18** | **0.17** |
| [EA_e]_0 | 16.40 | 0.32 | 16.01 | 0.35 | 16.00 | 0.46 | **16.14** | **0.23** |
| [GLY_e]_0 | 33.83 | 0.33 | 33.40 | 0.36 | 33.08 | 0.48 | **33.43** | **0.38** |
| [BIOMASS]_0 | 0.017 | 0.001 | 0.015 | 0.001 | 0.015 | 0.001 | **0.016** | **0.01** |
| [Ace_e_0]_0 | 0.30 | 0.16 | 0.28 | 0.17 | 0.26 | 0.23 | **0.28** | **0.02** |
| [Ace_e_1]_0 | 0.11 | 0.16 | 0.13 | 0.17 | 0.10 | 0.23 | **0.11** | **0.02** |
| [NH4_e]_0 | 0.30 | 0.31 | 0.48 | 0.34 | 0.33 | 0.46 | **0.37** | **0.10** |
| $v_{21}$ | 6.24 | 0.51 | 6.22 | 0.63 | 6.08 | 0.78 | **6.18** | **0.09** |

[a]best fit obtained with COPASI

[b]standard deviation estimated by COPASI from the best fit

[c]mean and standard deviation of the three independent biological replicates
