## Supplementary Data 3 for "Integrative *in vivo* analysis of the ethanolamine utilization bacterial microcompartment in *Escherichia coli.*"

-

Supplementary Data 3

Denis Jallet[1#], Vanessa Soldan[2], Ramteen Shayan[2], Alexandre Stella[3,5], Nour Ismail[1], Rania Zenati[1], Edern Cahoreau[1,4], Odile Burlet-Schiltz[3,5], Stéphanie Balor[2], Pierre Millard[1,4], Stéphanie Heux[1].

[1] Toulouse Biotechnology Institute, Université de Toulouse, CNRS, INRAE, INSA, Toulouse, France

[2] Plateforme de Microscopie Electronique Intégrative, Centre de Biologie Intégrative, Université de Toulouse, CNRS, Toulouse, France

[3] Institut de Pharmacologie et de Biologie Structurale (IPBS), Université de Toulouse, CNRS, Université Toulouse III – Paul Sabatier (UT3), Toulouse, France

[4] MetaToul-MetaboHUB, National infrastructure of metabolomics and fluxomics, Toulouse, France

[5] Infrastructure nationale de protéomique, ProFI, FR 2048, Toulouse, France

### Correspondance:

Here, the flux calculations through individual BMCs based on data from literature and our experimental results are described.

#### Flux through individual carboxysomes :

**A) Based on data from Reinhold et al., 1991 (10.1139/b91-126).**

*First calculation: data from paragraph 4 on page 986*

The authors give the following $V_{max}$ value for $CO_2$ fixation:

$V_{max}$ = $4.0 \times 10^{-5}$ [mol/ (cm$^3$ carboxysome s)]

Let the radius of a BMC be

$r_{BMC}$ = $2 \times 10^{-5}$ [cm]

Let the volume of a carboxysome be (NB: assuming a spherical shape for simplicity)

$v_{BMC}$ = $4/3 \times \pi \times r^3$

= $3.35 \times 10^{-14}$ [cm$^3$]

Vmax for a single BMC will be

$V_{max})_{BMC}$ = $V_{max} \times v_{BMC}$ [mol/(carboxysome s)]

= $1.34 \times 10^{-18}$ [mol/(carboxysome s)] or $1.34 \times 10^{-3}$ [fmol/(carboxysome s)]

= $4.82 \times 10^{-15}$ [mol/( carboxysome h)] or 4.82 [fmol/( carboxysome h)]

*Second calculation: data from paragraph 2 on page 987*

The authors mention the following experimental $V_{max}$ value for $CO_2$ fixation (derived from the work by Price and Badger 1989 (10.1139/b91-124)):

$V_{max\cdot exp})_{cell}$ = $2.6 \times 10^{-18}$ [mol/ (cell s)]

= $9.36 \times 10^{-15}$ [mol/ (cell h)]

Let's assume there are $N_{BMC})_{cell}$ = 6 carboxysomes per bacterium as mentioned by the authors in their text.

The $V_{max}$ on a per carboxysome basis will be:

$V_{max\cdot exp})_{BMC}$ = $V_{max\cdot exp})_{cell}$ / $N_{BMC})_{cell}$

= $4.33 \times 10^{-19}$ [mol/ (carboxysome s)] or $4.33 \times 10^{-4}$ [fmol/ (carboxysome s)]

= $1.56 \times 10^{-15}$ [mol/ (carboxysome h)] or 1.56 [fmol/ (carboxysome h)]

**B) Based on data from Mangan and Brenner, 2014 (10.7554/eLife.02043).**

Table 3: Carboxylation rate $8.2 \times 10^{-8}$ [picomoles/(cell s)] assuming a high $HCO_3^-$ transport rate resulting in 30 mM $HCO_3^-$ cytosolic pool.

$V)_{cell}$ = $8.2 \times 10^{-8}$ [picomol/(cell s)]

= $8.2 \times 10^{-8} \times 10^{-12} \times 3.6 \times 10^{3}$ [mol/(cell h)]

= $2.95 \times 10^{-16}$ [mol/(cell h)] or 0.295 [fmol/(cell h)]

Let's assume there are $N_{BMC})_{cell}$ = 6 carboxysomes per bacterium (NB: same assumption as in Reinhold et al., 1991).

$V)_{BMC}$ = $V)_{cell}$ / $N_{BMC})_{cell}$

= $4.9 \times 10^{-17}$ [mol/(carboxysome h)] or 0.049 [fmol/(carboxysome h)]

= $1.33 \times 10^{-20}$ [mol/(carboxysome s)] or $1.33 \times 10^{-5}$ [fmol/(carboxysome s)]

Table 4: Carboxylation rate $5.4 \times 10^{-8}$ [picomol/(cell s)] assuming a lower $HCO_3^-$ transport rate.

$V)_{cell}$ = $1.94 \times 10^{-16}$ [mol/(cell h)] or 0.194 [fmol/(cell h)]

$V)_{BMC}$ = $3.2 \times 10^{-17}$ [mol/(carboxysome h)] or 0.032 [fmol/(carboxysome h)]

#### Flux through individual Pdu BMCs :

**A) Work by Jakobson et al., 2017 (10.1371/journal.pcbi.1005525)**

Assuming that PduP/Q limit 1,2-PD assimilation within the *S. enterica* Pdu BMCs *in vivo*, the authors calculated the following 1,2-PD conversion flux per cell:

$F_{1,2\text{-}PD})_{cell}$ = $2.97 \times 10^{-13}$ [µmol/(cell s)]

= $1.07 \times 10^{-9}$ [µmol/(cell h)]

= $1.07 \times 10^{-15}$ [mol/(cell h)] or 1.07 [fmol/(cell h)]

Let assume the number of Pdu BMCs per cell to be (based on Kennedy et al., 2022; 10.1128/jb.00576-21)

$N_{BMCs})_{cell}$ = 3

The 1,2-PD conversion flux per Pdu BMC will then be

$V_{1,2\text{-}PD})_{BMC}$ = $F_{cell}$ / $N_{BMCs}$

= 1.07 / 3 = 0.356 [fmol/(BMC h)]

**B) Work by Jakobson et al., 2018 (10.1038/s41598-018-26399-0)**

For 1,2-PD assimilation, pathway flux of roughly

$F_{1,2\text{-}PD})_{cell}$ = $2.97 \times 10^{-13}$ [µmol/(cell s)] (based on observations by Sampson and Bobik, 2008: 10.1128/JB.01925-07)

The results will be identical to that above.

#### Flux through individual Eut BMCs :

Based on our experimental data, let the EA uptake rate be

$q_S)_{EA}$ = 7 [$mmol.g_{DW}^{-1}.h^{-1}$]

Let the number of Eut BMCs per cell be

$N_{BMCs})_{cell}$ = 6

Based on literature, let the dry weight of an *E. coli* cell be

$m_{cell}$ = 258 [fg] (BioNumbers assuming a doubling rate of 60 min)

= $2.58 \times 10^{-13}$ [g]

The number of *E. coli* cells in 1 g DW will be

$N_{cells}$ = $1 / m_{cell}$

= $1 / (2.58 \times 10^{-13}) = 3.88 \times 10^{12}$ cells

The number of Eut BMCs in 1 g DW will be

$N_{BMCs}$ = $N_{cells} \times N_{BMCs})_{cell}$

= $3.88 \times 10^{12} \times 6 = 2.33 \times 10^{13}$

The EA conversion flux per BMC will be

$V_{EA})_{BMC}$ = $q_{S})_{EA} / N_{BMCs}$

= $7 / (2.33 \times 10^{13}) = 3.00 \times 10^{-13}$ [mmol/(BMC h)]

= $3.00 \times 10^{-16}$ [mol/(BMC h)]

= 0.300 [fmol/(BMC h)]

**Glycolytic flux based on our data:**

Based on our modelling data, let the glycolytic flux be:

$F_{glyco}$ = 2.5 [$mmol.g_{DW}^{-1}.h^{-1}$]

Based on literature, let the dry weight of an *E. coli* cell be

$m_{cell}$ = 258 [fg] (BioNumbers assuming a doubling rate of 60 min)

= $2.58 \times 10^{-13}$ [g]

The number of *E. coli* cells in 1 g DW will be

$N_{cells}$ = $1 / m_{cell}$

= $1 / (2.58 \times 10^{-13}) = 3.88 \times 10^{12}$ cells

The glycolytic flux per cell will be:

$F_{glyco)cell}$ = $F_{glyco} / N_{cells}$

= $2.5 / (3.88 \times 10^{12})$ [mmol/(cell h)]

= $7.2 \times 10^{-13}$ [mmol/(cell h)]

= $7.2 \times 10^{-16}$ [mol/(cell h)]

= 0.72 [fmol/(cell h)]

PEP step)

Then $F_{glyco)cell}$ = 2.016 [fmol/(cell h)]
